# Developmental Genome-Wide DNA Methylation Asymmetry Between Mouse Placenta and Embryo

**DOI:** 10.1101/718247

**Authors:** LM Legault, K Doiron, A Lemieux, M Caron, D Chan, FL Lopes, G Bourque, D Sinnett, S McGraw

## Abstract

In early embryos, DNA methylation is remodelled to initiate the developmental program but for mostly unknown reasons, methylation marks are acquired unequally between embryonic and placental cells. To better understand this, we generated high-resolution DNA methylation maps of mouse mid-gestation (E10.5) embryo and placenta. We uncovered specific subtypes of differentially methylated regions (DMRs) that contribute directly to the developmental asymmetry existing between mid-gestation embryonic and placental DNA methylation patterns. We show that the asymmetry occurs rapidly during the acquisition of marks in the post-implanted conceptus (E3.5-E6.5), and that these patterns are long-lasting across subtypes of DMRs throughout prenatal development and in somatic tissues. We reveal that at the peri-implantation stages, the *de novo* methyltransferase activity of DNMT3B is the main driver of methylation marks on asymmetric DMRs, and that DNMT3B can largely compensate for lack of DNMT3A in the epiblast and extraembryonic ectoderm, whereas DNMT3A can only partially compensate in the absence of DNMT3B. However, as development progresses and as DNMT3A becomes the principal *de novo* methyltransferase, the compensatory DNA methylation mechanism of DNMT3B on DMRs becomes less effective.

## INTRODUCTION

Throughout the eutherian mammalian gestation, the placenta plays an essential role in mediating maternal–embryonic exchanges of gas, nutrients and waste, and also provides the developing embryo with a protective layer against adverse environmental exposures and the maternal immune system (Rossant & Cross, 2001). These unique placental functions are orchestrated by several distinct trophoblast cell subtypes organized in separate layers (Cross, 2000). The initial steps of lineage specialization of both placental and embryonic cells occur promptly following fertilization during the first few embryonic cleavages as DNA methylation marks are being reprogrammed (Morgan, Santos, Green, Dean, & Reik, 2005).

DNA methylation is an epigenetic mechanism that is critical in the determination of lineage-specific differentiation and development, and is mainly recognized for its involvement in processes such as transcriptional repression, genomic imprinting and X-inactivation (Bestor, 2000). DNA methylation marks are mediated by the action of DNA methyltransferases (DNMTs). Establishment of new or *de novo* DNA methylation patterns required for cell lineage determination during development is mediated by DNMT3A and DNMT3B, with cofactor DNMT3L, (Li, 2002; Okano, Bell, Haber, & Li, 1999), whereas DNMT1 maintains heritable DNA methylation patterns during cellular divisions (Lei et al., 1996; Leonhardt, Page, Weier, & Bestor, 1992). These enzymes are critical, as deletion of *Dnmt3b* or *Dnmt1* is embryonic lethal, while *Dnmt3a*-deficient offspring die shortly after birth (Li, Bestor, & Jaenisch, 1992; Okano et al., 1999). During gametogenesis, the acquisition of genome-wide and allele-specific methylation patterns (i.e. genomic imprinting) in both oocytes and sperm is essentially due to the activity of DNMT3A (Kaneda et al., 2004; Kato et al., 2007). Following fertilization, a reprogramming wave removes most methylation signatures across the genome, except for imprinted regions, some types of repeat sequences, as well as imprinted-like sequences, to trigger the developmental program (Hirasawa et al., 2008; Howell et al., 2001; McGraw et al., 2015). Then, during the peri-implantation process, DNA methylation profiles are re-acquired in a sex-, cell- and tissue-specific manner across most parts of the genome by the combined action of DNMT3A and DNMT3B. In the early stages of the *de novo* methylation wave (E4.5-E7.5), the expression of *Dnmt3b* is more robust than *Dnmt3a* in the epiblast and embryonic-derived cells (Auclair, Guibert, Bender, & Weber, 2014; Smith et al., 2017; Watanabe, Suetake, Tada, & Tajima, 2002), with the relative expression of *Dnmt3b* and *Dnmt3a* being considerably reduced in the extraembryonic ectoderm (E×E) and trophoblast lineages (Senner, Krueger, Oxley, Andrews, & Hemberger, 2012; Smith et al., 2017). This discrepancy in *Dnmt3a* and *Dnmt3b* expression levels coincides with the initiation of divergent DNA methylation acquisition between the trophoblast and the inner cell mass of the blastocyst (Fulka, Mrazek, Tepla, & Fulka, 2004; Guo et al., 2014; Monk, 1987; Nakanishi et al., 2012; Oda, Oxley, Dean, & Reik, 2013; Santos, Hendrich, Reik, & Dean, 2002; Smith et al., 2014), a difference that becomes extremely apparent by E6.5, as the epiblast has acquired most of its global DNA methylation compared to the lower-methylation state of the E×E (Auclair et al., 2014; Smith et al., 2017; Zhang et al., 2018). This divergence is a common feature across mammalian placenta, as a heterogeneous and lower-methylation state compared to somatic tissues and other cell types is constantly observed (Chatterjee et al., 2016; Decato, Lopez-Tello, Sferruzzi-Perri, Smith, & Dean, 2017; Schroeder et al., 2013; Smith et al., 2017).

Although the functional role of reduced methylation levels observed across the placental genome is still not fully understood, studies suggest that it may activate transposable elements that are typically silenced in other tissues (Chuong, 2013). DNA methylation plays an important role in suppressing retrotransposons in mammalian cells, for which the activity has been associated with genomic instability and disease development (Church et al., 2009; Slotkin & Martienssen, 2007). Following the *de novo* methylation wave, in embryonic-derived cells from the inner cell mass, transposable elements acquire higher levels of DNA methylation causing transcriptional silencing, whereas in the trophectoderm-derived cells that will form the placenta, these transposable elements are maintained in a relaxed methylation state and preferentially expressed (Okahara et al., 2004; Price et al., 2012; Warren et al., 2015). The low-methylation levels on these elements contributed to the evolution and diversification of the placenta function through the regulation of gene expression by providing placenta-specific enhancers, cryptic promoters and other cis-regulatory elements (Cohen et al., 2011; Emera & Wagner, 2012; Haig, 2012; Macaulay, Weeks, Andrews, & Morison, 2011; Mi et al., 2000; Xie et al., 2013).

Despite the evident distinction between embryonic and placental DNA methylation levels, it remains unclear how, when and where methylation levels are acquired unequally across these genomes. To better understand the developmental dynamics of epigenetic asymmetry that exists between embryo and placenta, we first generated high-resolution maps of DNA methylation marks using Reduced Representation Bisulfite Sequencing (RRBS) at mid-gestation, when the mouse placenta is first considered mature (E10.5), to identify embryo-placenta differentially methylated regions (DMRs). Then, using publicly available DNA methylation data sets and computational analyses, we defined how various categories of DMRs are established in early stages and maintained throughout development. In addition, we outlined the contribution of *Dnmt3a* and *Dnmt3b* in the acquisition and maintenance of embryonic and extraembryonic specific DMR patterns.

## METHODS

### Animals and Sample Collection

Female C57BL/6N (8-10 week-old) were purchased from Harlan Sprague-Dawley Laboratories (Indianapolis, IN) and were mated with male C57BL/6N (4 months of age). Following natural mating, embryos and placental tissues were collected at E10.5 (presence of vaginal plug at E0.5). Maternal decidua was removed from placentas. Samples were frozen immediately in liquid nitrogen and stored at −80 °C until analyzed.

### DNA Methylation Analyses

RRBS libraries were generated as published protocols (Boyle et al., 2012; Gu et al., 2011) with our specifications (Legault, Chan, & McGraw, 2019; Magnus et al., 2014; McGraw et al., 2015). 500 ng of extracted DNA (Qiagen) from placenta (male n=2, female n=2) and embryo (male n=2, female n=2) samples was *MspI* digested, adaptor ligated and PCR amplified (multiplex). Multiplexed samples were pooled and 100 bp paired-end sequenced (HiSeq-2000, Illumina). The data analyses were done according to the pipeline established at the McGill Epigenomics Mapping and Data Coordinating Centers (Magnus et al., 2014; McGraw et al., 2015) that include BSMAP and methylKit. Specific parameters were chosen including 100 bp step-wise tiling windows, containing a minimum of 2 CpGs per tile and a minimum 15× CpG coverage of each tile per sample. The methylation level of a 100-bp tile was the result of all CpG C/T read counts within the tile after coverage normalization between samples, and the methylation level reported for a sample on autosomal chromosomes was the average methylation level across all individual replicates. Significant DNA methylation changes were designated as ±≥20% average differences between groups of replicates and a q-value < 0.01 using the logistic regression function of methylKit (Akalin et al., 2012). Direct comparisons between DNA methylation averages were done using Wilcoxon-Mann-Whitney test in R. Gene ontology (GO) terms and pathway analyses for RefSeqs associated to promoter-TSS were conducted using Metascape gene annotation and analysis resource (Zhou et al., 2019).

### Sequencing Data

Publicly available DNA methylation data sets (Auclair et al., 2014; Decato et al., 2017; Hon et al., 2013; Smith et al., 2014; Smith et al., 2017; Whidden et al., 2016) were analyzed using a custom script to intersect single CpG site methylation calls from these data sets within defined 100bp tiles associated to the embryo-placenta DMRs categories

## RESULTS

### Increased Fluctuation in Genome-Wide DNA Methylation Levels in Placental Cells

To identify the overall epigenetic asymmetry that exists between placental and embryonic genomes during mouse *in utero* development, we first established genome-wide DNA methylation profiles using RRBS (Legault et al., 2019; Magnus et al., 2014; McGraw et al., 2015) of embryos and their corresponding placentas at mid-gestation (E10.5), the developmental stage at which the mouse placenta is considered mature (Cross, Werb, & Fisher, 1994). Using this approach, we quantified the DNA methylation profiles of ~1.8 million CpG sites in each sample (embryo n=4, placenta n=4). We found that the accumulation of CpG methylation was very distinct between the placenta and the embryo. In the placenta, we detected a greater proportion of CpGs in the 0-50% methylation range and a lower proportion of CpGs in the 80-100% methylation range (Figure 1A). Although we observed that a large proportion of CpGs within the examined regions had no methylation marks in both the embryo and the placenta, interestingly, most CpGs with partial methylation (20-50%) in placenta showed high methylation levels (>80%) in embryos (Figure 1B). The divergence in global DNA methylation profiles conferred a high degree of clustering between both tissue types (Figure S1). These results are consistent with previous studies indicating lower levels of overall DNA methylation in extraembryonic tissues (Price et al., 2012; Schroeder et al., 2013; Smith et al., 2017), reviewed in (Robinson & Price, 2015).

**Figure 1.**
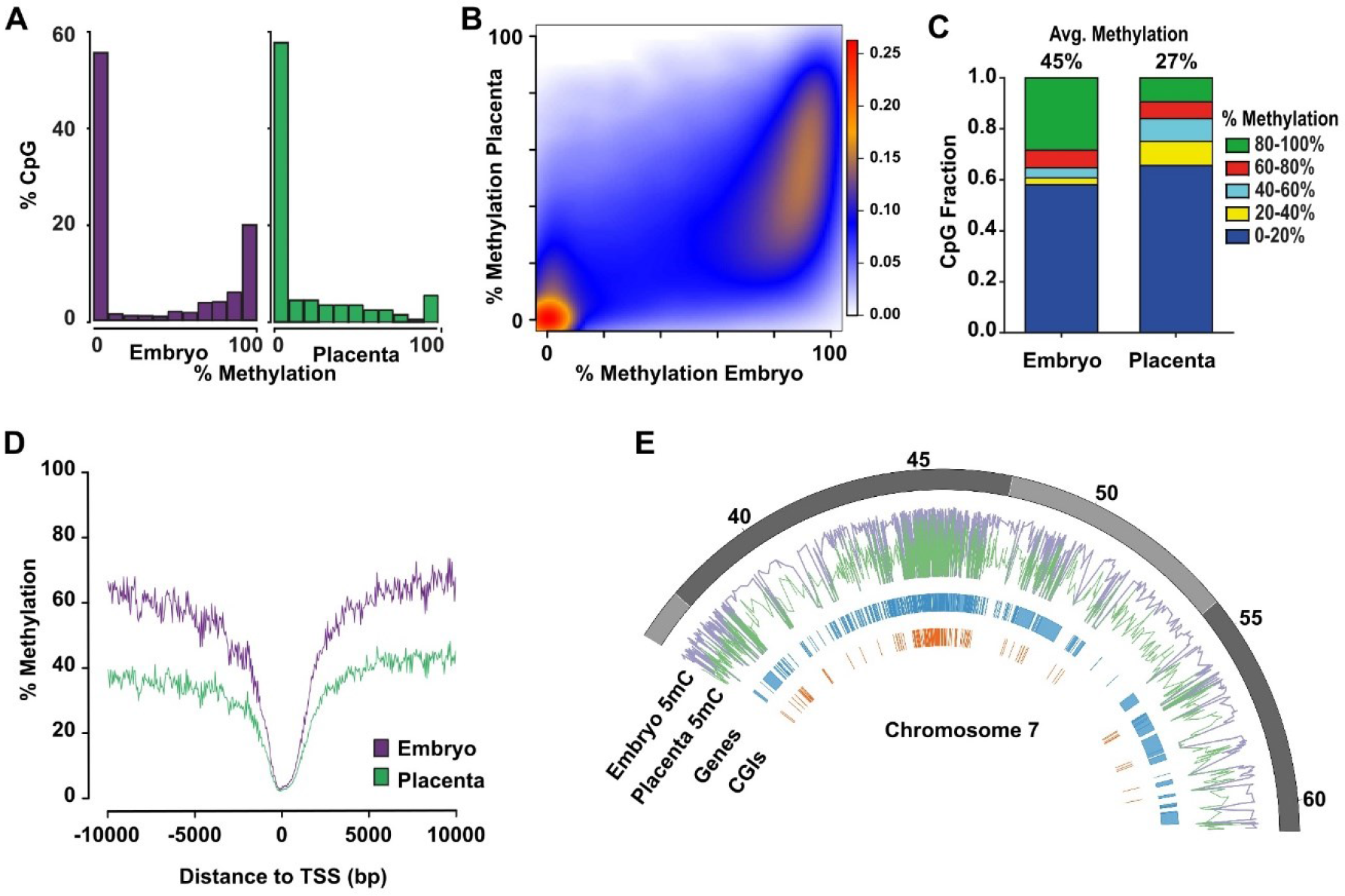
Distinctive patterns of genome-wide DNA methylation accumulation between E10.5 embryo and placenta. Analyses of genome-wide DNA methylation sequencing results for embryo (n=4) and placenta (n=4). **A)** Density histograms showing the distribution of CpG methylation levels for embryo (*purple*) and placenta (*green*). **B)** Pairwise comparison of CpG methylation between embryo and placenta. Density increases from blue to red. **C)** CpG fraction of 100 bp tiles within 0-20%, 20-40%, 40-60%, 60-80% and 80-100% ranges in embryo and placenta. Shown above bars, is the average CpG methylation for each unique tile represented in graph. **D)** DNA methylation means surrounding the transcription start site (TSS) for All-tiles in each experimental group. **E)** Circle plot showing methylation average of embryo (*purple*) and placenta (*green*) across a portion of chromosome 7.

To further investigate the dynamics of DNA methylation between placental and embryonic genomes and enable direct comparison of precise regions, we segmented the genome of autosomal chromosomes into 100bp non-overlapping genomic windows (*tiles*; see methods section). After removal of sex chromosomes, we identified 245 048 unique sequenced tiles (referred to as All-tiles) containing 896 820 common CpGs between all placenta and embryo samples and with a minimum of 15× sequencing depth. We observed a strong reduction in the fraction of highly methylated tiles (80-100%) in the placenta compared to the embryo (Figure 1C), which correlated with a sharp increase in the number of placenta tiles in the 0-20%, 20-40% and 40-60% methylation range. Globally, we found that the average DNA methylation level across all placental tiles was significantly lower compared to all embryo tiles (27% vs 45%, p<0.0001) (Figure 1C). This overall epigenetic disparity in embryo and placenta DNA methylation levels was especially noticeable when we mapped the methylation mean of All-tiles with respect to regions surrounding the transcription start sites (TSS) (Figure 1D). When we focused on a specific chromosome section (e.g., chr 7, 35Mb) (Figure 1E), we observed that genomic segments with high DNA methylation levels in embryos had predominantly lower levels in placenta. We also observed that gene or CpG island (CGI) poor regions had consistently high methylation levels in embryos and lower methylation levels in the placenta. Together, these results indicate that the mid-gestation placenta has very distinctive global DNA methylation levels compared to the embryo, and that this lower level of global DNA methylation across the placental genome is due to a significant lower number of highly methylated (≥80%) genomic regions. We can also conclude that despite the cellular heterogeneity in E10.5 placental and embryonic tissues, the vast majority of the conceptus possesses specific genomic regions with either low (0-20%) or high (80-100%) levels of methylation. However, the placenta genome presents an increased number of regions having a broader distribution of DNA methylation (20-80%), revealing a greater diversity in methylation levels across placental cell types compared to embryonic cell types.

### Embryonic and Placental DNA Methylation Divergences across Genomic Features

To explain the developmentally divergent methylated states between the mouse embryo and the placenta, we next sought to precisely determine the genomic features revealing DNA methylation differences. We defined differentially methylated regions (DMRs) as 100bp genomic segments showing a significant difference of methylation levels between embryonic and placental samples with an absolute mean methylation difference of 20% or higher (McGraw et al., 2015; Piche et al., 2019; Shaffer et al., 2015). Using these conditions, we screened the 245 048 unique tiles common between all samples and identified 110 240 DMRs (~45% of All-tiles; Supplementary Table 1) with tissue- and/or developmental-specific DNA methylation variations between the embryo and the placenta (Figure2A; random subset of 20,000 DMRs shown). Consistent with our findings (Figure 1), the majority (96.8%; n=106 712) of DMRs had lower DNA methylation in the placenta (referred to as Hypo-DMRs) and only a small proportion (3.2%; n=3 528) showed increased methylation levels (referred to as Hyper-DMRs) compared to the embryo (Figure S2). For Hypo-DMRs, tiles mainly overlapped (96%; n=102 982) with intergenic, intron, exon and promoter-TSS regions (Figure S2). For each of the genomic feature categories of Hypo-DMRs, the average DNA methylation levels in the placenta were essentially half of those present in the embryo (Figure 2B). As for DMRs with higher methylation levels in the placenta (Hyper-DMRs), we noticed that the vast majority of these tiles (83%; n=2 929) had low methylation levels in the embryo (<20%) (Figure 2A-B). Most of Hyper-DMRs overlapped intergenic, intron, exon and promoter-TSS regions (93%; n=3 272, Figure S2). We then investigated the relationship between gene expression and DMR DNA methylation levels. Highly expressed genes in placental tissue such as *Syna* (*Syncytin A*) (Mi et al., 2000) showed overall lower methylation levels in the placenta when compared to the embryo (Figure 2D, smoothed representation) (Hansen, Langmead, & Irizarry, 2012). Similar observations were made in the gene body of *Atf6b* (*Activating Transcription Factor 6 Beta*), a gene implicated in the transcriptional downregulation of *Pgf* (*Placental Growth Factor*) in response to endoplasmic reticulum stress in pathological placentas (Mizuuchi et al., 2016). As for genes predominantly expressed in the embryo, *Mir219a-2/Mir219b*, a brain-specific non-coding microRNA, showed higher methylation in the placenta (Figure 2D). It was recently shown that *Mir219* was abnormally expressed during pregnancy in amnion retrieved specifically from obese women (Nardelli et al., 2014) and, interestingly, obesity has been suggested to cause epigenetic changes (Lillycrop & Burdge, 2011). Another example of placenta Hyper-DMRs is *Sox6* (*SRY-Box 6*), which is implicated in the terminal differentiation of muscle (Figure 2D) (Kamachi & Kondoh, 2013). These observations point to an association between tissue-specific DNA methylation levels of DMRs and biological functions. Gene ontology enrichment analyses further support that promoter-associated Hypo-DMRs were strongly associated to germline functions and reproduction (*e.g.*, male and female gamete generation, reproduction, piRNA metabolic process, meiotic cell cycle, germ cell development) (Figure 2C). As for promoter-associated Hyper-DMRs, top biological processes were mostly associated with developmental and differentiation processes (*e.g.*, regionalization, embryo development, pattern specification process, skeletal system, head development) (Figure 2C). Fittingly, the biological functions were completely divergent between Hypo- and Hyper-DMRs. Altogether, these results denote that Hypo- and Hyper-DMRs are present across genomic features between mid-gestation embryo and placenta, and that these DNA methylation divergences are implicated in promoting/repressing specific processes during embryonic and placental development.

**Figure 2.**
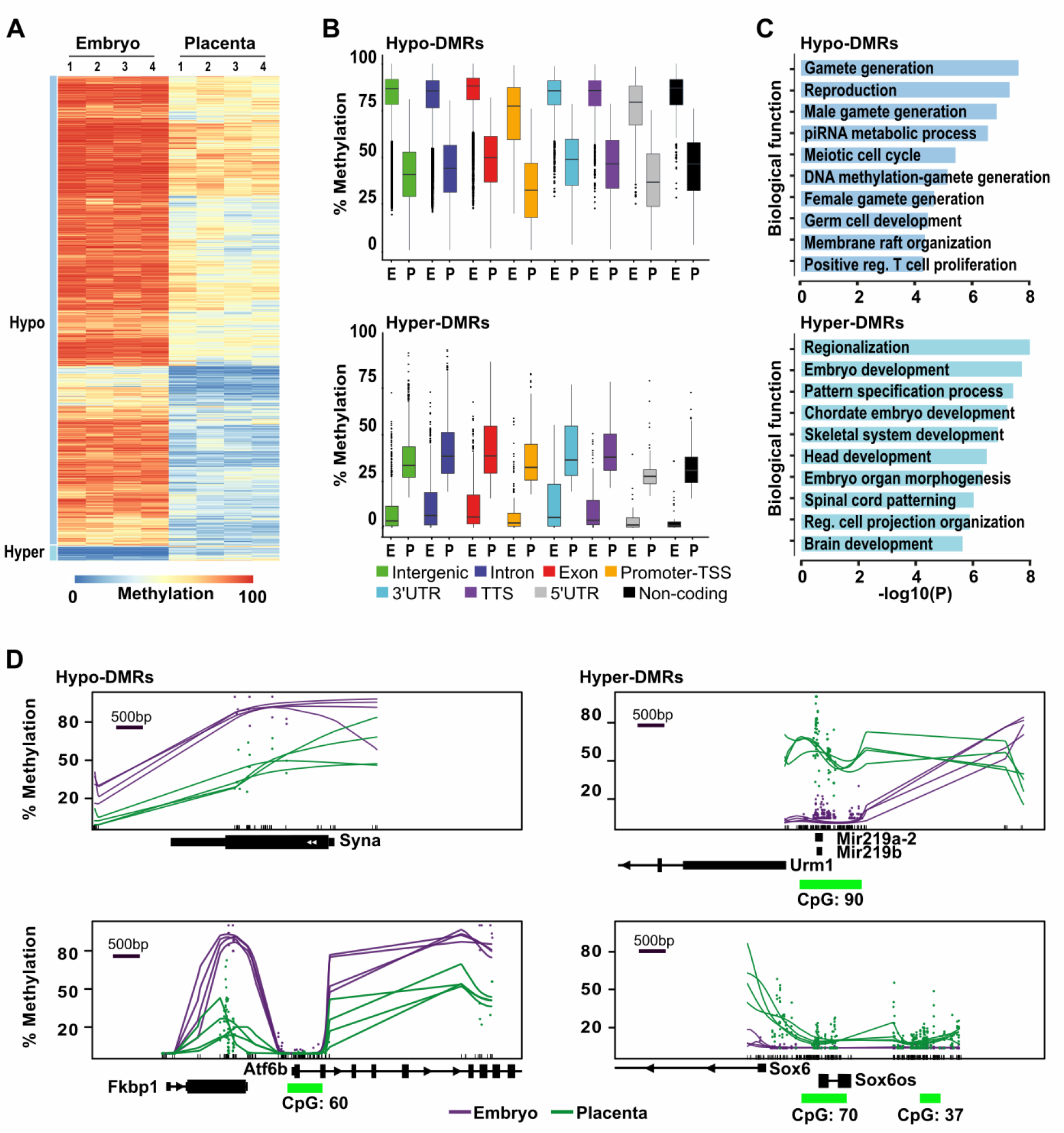
Genome-wide methylation asymmetry across genomic features between E10.5 embryo and placenta. Analyses of genome-wide DNA methylation sequencing results for embryo (n=4) and placenta (n=4) samples. **A)** Heatmap representation of DNA methylation levels for the 20 000 tiles with the most variable levels between embryo and placenta (DNA methylation variance >20% and *P*<0.05). **B)** Box-plots representing DNA methylation distribution in embryo and placenta for the different genomic annotation regions (intergenic, intron, exon, promoter-TSS, 3’UTR, TTS, 5’UTR and non-coding). **C)** Summary of biological functions associated with promoter regions in Hypo- and Hyper-DMRs. **D)** Examples of smoothed methylation profiles (*BSmooth* tool) in regions with lower methylation profiles (*Syna and Atf6b*) or higher methylation profiles (*Mir219a-2/Mir219b*, and *Sox6*) in placenta.

### Presence of Distinctive DMR Categories between Mid-Gestation Embryo and Placenta

Amongst DMRs, our analyses also suggest the presence of particular DMR categories based on their level of DNA methylation in the embryo and placenta (Figure 2A). By defining subsets of DMRs and establishing their dynamic properties between tissues, we might better understand the genome-wide asymmetry in DNA methylation levels observed between the embryo and the placenta. To do so, we first clustered DMR-associated tiles in 6 different categories based on their range of low, mid and high methylation level (Low; <20%, Mid; ≥20 to <80%, High; ≥80%) in the embryo and the placenta, and followed the DMR category transitions between both tissues. We observed that DMRs with High-levels of methylation in the embryo overlapped with a large proportion of DMRs that showed Mid-levels of methylation in the placenta (Figure 3A,B; High-Mid n=72 715), whereas only a fraction corresponded to DMRs with Low-levels of methylation in the placenta (Figure 3A,B; High-Low n=1 889). As for DMRs with Mid-levels of methylation in embryo, the largest part remained in that same Mid-levels category in the placenta (Figure 3A; Mid-Mid n=23 640). Nonetheless, a portion of these embryonic Mid-levels DMRs were directed to DMRs with either Low- (Figure 3A; Mid-Low n=9 055) or High- (Figure 3A; Mid-High n= 21) levels of methylation in the placenta. Finally, DMRs with Low-levels of methylation in the embryo all showed Mid-levels of methylation in the placenta (Figure 3A; Low-Mid n=2 920). Thus, when we subdivide our DMRs into distinctive categories, we uncover that embryonic cells possess a large proportion of DMRs of High-level (≥80%) of DNA methylation, which remain potentially static in embryonic tissues, whereas the vast majority of these DMRs have lower and wide-range DNA methylation levels within placental tissue.

**Figure 3.**
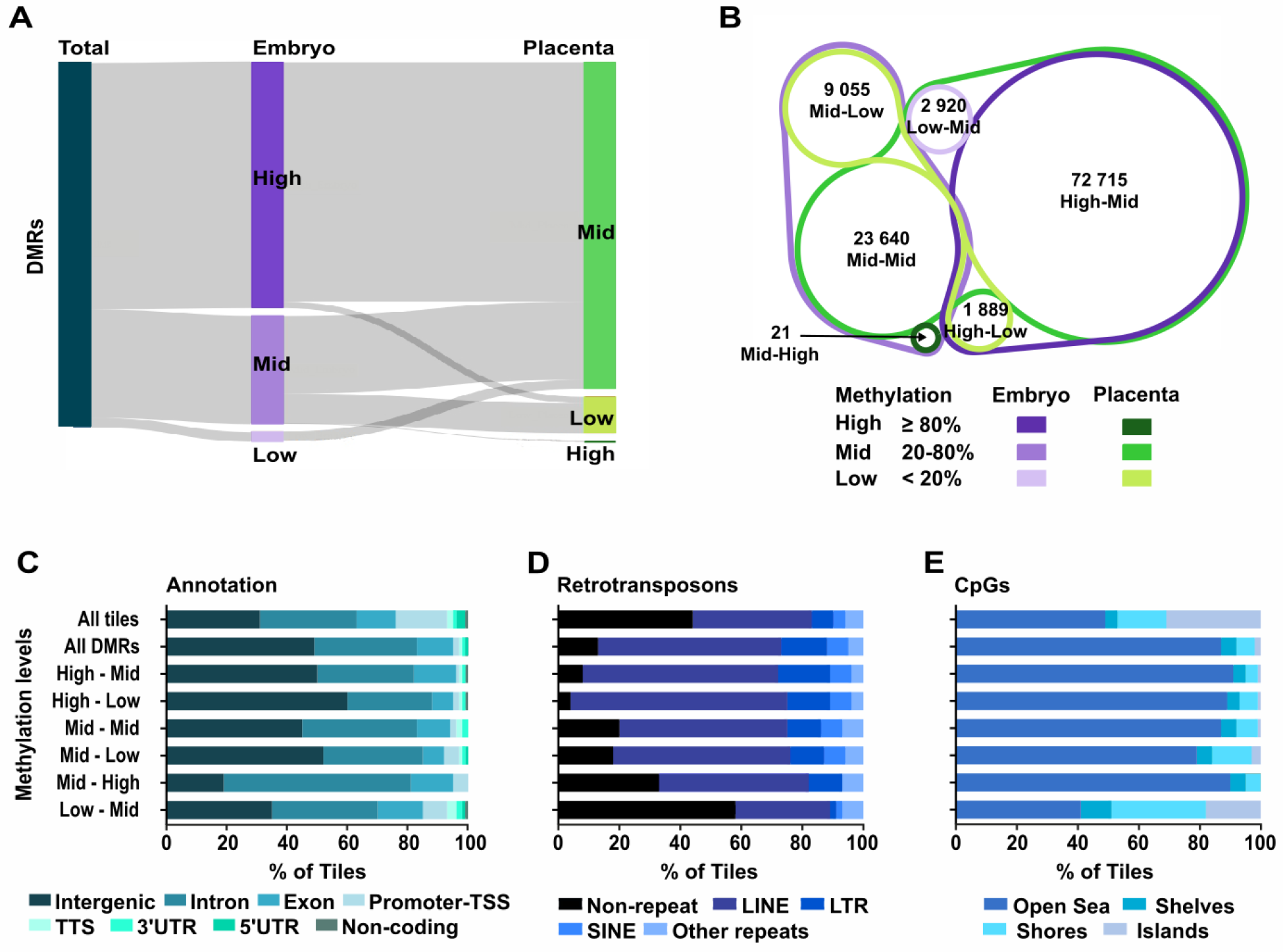
Distinct genomic features in DMRs based on their methylation status in E10.5 embryo and placenta. DMR analysis by methylation levels in embryo and placenta. **A)** Sankey diagram dividing DMRs by methylation levels in embryo (*purple*) and placenta (*green*) for the associated tiles. High : ≥80% methylation, Mid (intermediate) : ≥20% - <80% methylation, Low : <20% methylation. **B)** Venn diagram showing the proportion of tiles in the different DMR categories based on DNA methylation levels between embryo and placenta. **C), D)** and **E)** Analysis of all tiles, all the DMRs, as well as the 6 different DMR categories based on levels of DNA methylation in embryo and placenta for : **C)** Genomic annotations, **D)** Main retrotransposons and **E)** Proximity of CpG rich regions. Neighboring CpG dense regions were defined as shore; up to 2kb away from CGIs, shelf; 2-4kb away from CGIs, and open sea; >4kb away from CGIs.

### Specific Genomic Features Associated with DMR Categories

We next aimed to determine if the genomic distribution of embryo-placenta DMR categories was associated with distinct genomic features. First, by classifying by genomic annotations, we observed that DMRs with reduced methylation levels in the placenta (High-Mid, High-Low, Mid-Low) were prevailingly found in intergenic regions (>50% of tiles) (Figure 3C), whereas DMR categories with equivalent (Mid-Mid) or greater methylation level (Mid-High, Low-Mid) in the placenta were more frequent in genic associated regions (>50% of tiles). However, divergence between DMR categories was observed when we performed ontology analyses on promoter regions (Figure S3), as each DMR subtype clearly showed distinct biological functions. For example, High-Mid DMRs top functions were related to genes (e.g., *Aszl, Tnp1*, *Piwil2*) associated with male gamete generation, piRNA metabolic process and male meiotic nuclear division, whereas those of the Low-Mid category were mostly associated with developmental aspects (e.g., skeletal system development, regionalization, nervous system development). Since we know that in placenta, activation of retrotransposon-derived genes is interrelated with low DNA methylation levels (Cohen et al., 2011; Macaulay et al., 2011; Reiss, Zhang, & Mager, 2007), we next assessed how DMR categories overlapped with major types of retrotransposons (LINE; long interspersed nuclear elements, SINE; short interspersed nuclear element, and LTR; long terminal repeats). Out of the DMR categories, those with High-levels of methylation in the embryo and either Mid- or Low-levels in the placenta (High-Mid and High-Low) showed the most enriched overlap with retrotransposons, with 92% and 96% respectively (Figure 3D). DMR categories associated with higher level of methylation in the placenta (Mid-High and Low-Mid) showed the least overlap with retrotransposons, especially the Low-Mid subtype. This highlights that during the *de novo* acquisition of DNA methylation patterns, DMRs with High-levels of methylation in the embryo and lower levels in the placenta are almost exclusively within retrotransposons-associated sequences, whereas DMRs with Low-levels of methylation in the embryo and higher DNA methylation in the placenta are preferentially outside retrotransposons-associated sequences. Finally, we investigated the proximity of the DMR categories in regards to CpG rich (CpG islands; CGI), neighbouring (shore; < 2kb away from CGIs, shelf; 2-4kb away from CGIs) and distant (open sea; > 4kb away from CGIs) regions (Figure 3E). We observed that most DMR categories are depleted from CGIs and are mostly found in open sea regions. In contrast, ~60% of tiles in Low-Mid DMRs overlapped with CGI, shores and shelves, revealing that the acquisition of *de novo* methylation for these genomic fragments in the extraembryonic lineage preferentially targets sequences inside or surrounding CGIs.

Altogether, these results indicate that the asymmetry within DMR categories, based on their methylation levels in the embryo and the placenta, can be associated to specific biological functions and genomic-derived features (e.g., CpG, retrotransposon contents). This is particularly apparent for DMRs with High-levels in the embryo (High-Mid, High-Low) and those with higher methylation levels in the placenta (Low-Mid).

### DMR Categories are Established During the De Novo Methylation Wave and Maintained Throughout Development

To gain insights into the kinetics of lineage-specific DMR establishment between mid-gestation embryo and placenta, as well as their status during development, we assessed the levels of methylation associated with tiles for each DMR category as a function of their developmental stage. Publicly available sequencing data (Smith et al., 2012; Whidden et al., 2016) were analyzed using our custom script to generate 100bp tiles, and the DNA methylation levels for each tile were calculated. In E3.5 blastocysts, when the mouse genome is mostly depleted from DNA methylation marks, we observed low global DNA methylation levels (average <20%) for all DMR subtypes, with similar median levels between the committed cells of the inner cell mass (ICM) and trophectoderm lineages (Figure 4). DNA methylation levels tended to be higher in the trophectoderm for regions falling in the Mid-High DMRs, although measurements are based on very few DMRs for this specific category (n=21, Figure 3B, Mid-High). Since global DNA methylation is re-acquired in the next few subsequent developmental stages, we then asked whether the contrast in DNA methylation levels associated with the various DMR categories at mid-gestation would already be present between E6.5 epiblast and extraembryonic ectoderm (E×E) cell lineages, layers that are mostly composed of homogeneous and undifferentiated cell populations. For all DMR categories, DNA methylation levels in the E6.5 epiblast and E×E already showed similar pattern trends to those observed in the E10.5 embryo and placenta (Figure 4, Table S2). Interestingly, for all DMR categories we observed slightly higher DNA methylation levels in E6.5 E×E compared to E10.5 placenta (Figure 4, Table S2). However, when measured in the subsequent developmental stages (E10.5, E11.5, E15 and E18), levels associated with each category stabilized and remained in the same methylation range. This is true for all categories, except for the Mid-High DMRs where methylation levels are reduced by ~20% at E18. As for the E6.5 epiblast DNA methylation profiles, they closely matched those observed in the E10.5 embryo, except again for the Mid-High DMRs. For each of the DMR subtypes, the DNA methylation levels were largely stable and persisted across other time points (E6.5, E10.5, E11.5) in embryonic cells. Although no public DNA methylation data were available for whole embryos at later stages, when we overlapped tiles associated with E10.5 DMR categories with data from differentiated tissues of adult mice, the global DNA methylation profiles closely matched those for the majority of DMR categories (High-Mid, High-Low, Mid-Mid and Mid-Low) (Figure 4, Figure S4). We conclude that the various DMR categories observed at E10.5 are established during the embryonic and extraembryonic lineage-specification processes occurring during the peri-implantation wave of *de novo* methylation, and that these DNA methylation landscapes are widely retained throughout embryo and placenta development. Furthermore, for most DMR categories, the DNA methylation levels observed in developing embryos are long-lasting and conserved throughout somatic cell differentiation.

**Figure 4.**
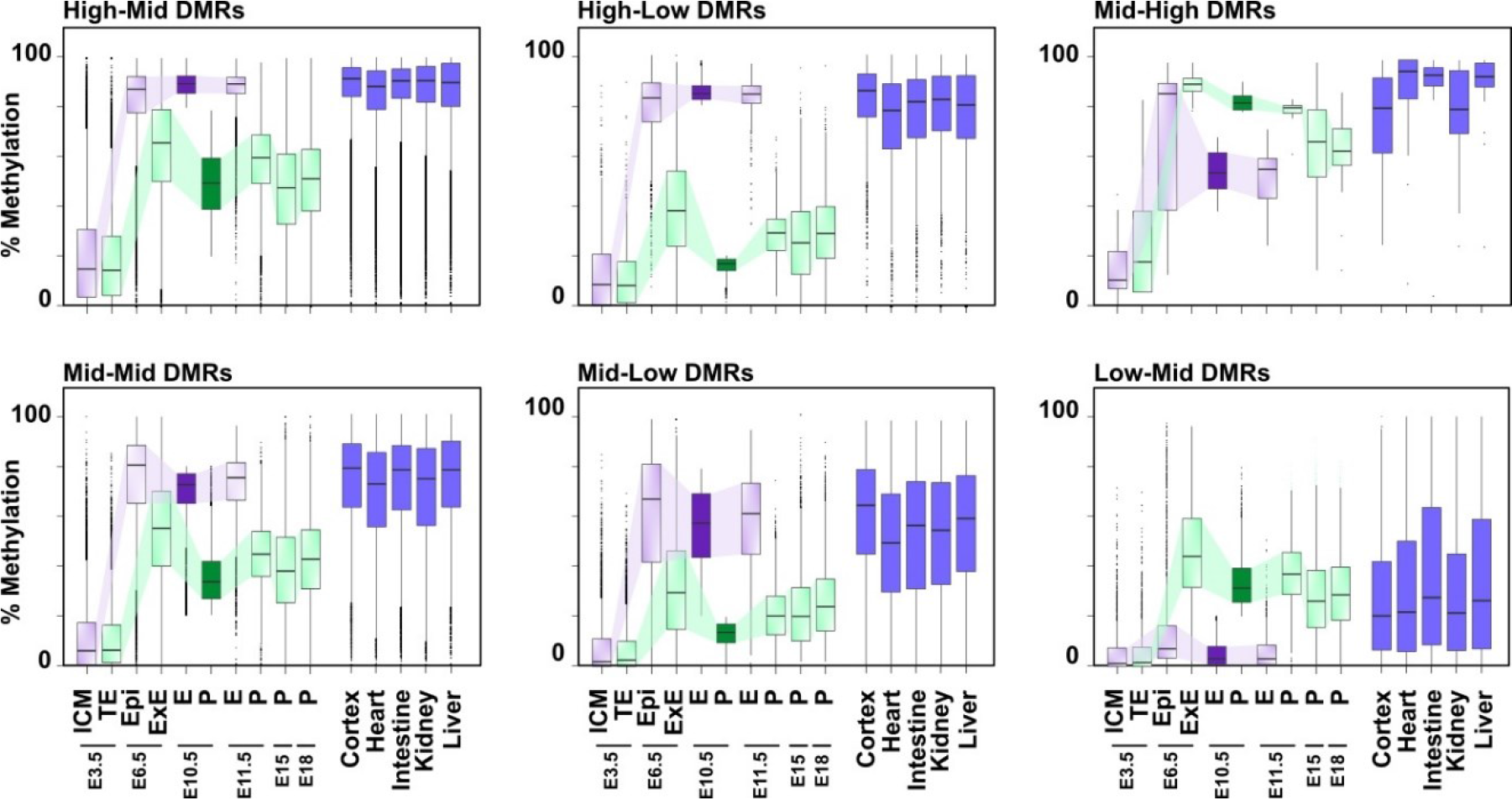
Dynamics of DNA methylation profiles associated with DMR categories and their evolution across embryo and placenta development. Box plots representing the DNA methylation distribution and median values for each DMR category in various developmental stages. Tiles associated with DMR categories at E10.5 were overlapped with previously published and publicly available data, and methylation levels were determined at each developmental stage (Smith et *al* 2012, Smith et *al.* 2014, Whidden et *al.* 2016) or in adult somatic tissues (Hon et *al.* 2013). ICM: inner cell mass, TE: trophectoderm, Epi: epiblast, E×E: extraembryonic ectoderm, E: embryo, Pla: placenta. See Table S2 for median and mean methylation values, and Table S4 for number of overlapping tiles analysed.

### Dnmt3a- or Dnmt3b-Deficiency Alters Proper Establishment of DMR-Associated Patterns

Since the combined activity of DNMT3A and DNMT3B is essential for proper establishment of DNA methylation profiles and normal development, we next sought to define the contribution of each enzyme in the *de novo* establishment of DNA methylation in subtypes of DMRs in embryonic and extraembryonic cell lineages. To do so, we used DNA methylation data from publicly available datasets (whole genome bisulfite sequencing and RRBS) at E6.5 (epiblast and E×E) (Smith et al., 2017) and E8.5 (embryo) (Auclair et al., 2014) with various inactive forms of *Dnmts*. First, when we overlapped our All-tiles subset, we observed a substantial reduction (p<0.0001) in average DNA methylation in absence of DNMT3A (37.81%) or DNMT3B (31.48%) in the E6.5 epiblast compared to wild-type (44.81%), whereas in the E6.5 E×E such a comparable loss was only associated with a *Dnmt3b-*deficiency (wt: 34.38% vs *Dnmt3b*^−/−^: 19.33%, p<0.0001) (Figure S5). Similarly, in the E8.5 embryo, a significant reduction in average DNA methylation level was measured with lack of *Dnmt3a* or *Dnmt3b* expression compared to wild-type (wt: 47.5%; *Dnmt3a*^−/−^: 43.6%; *Dnmt3b*^−/−^: 34%, p<0.0001) (Figure S5). For most DMR subtypes, absence of DNMT3A caused modest or no reduction on overall DNA methylation levels in E6.5 epiblast and E×E (Figure 5A, Table S3). Intriguingly, *Dnmt3a-*deficiency caused a slight gain of methylation (wt: 76.9% vs *Dnmt3a*^−/−^: 81.5%, p<0.0001) level in the E6.5 epiblast for High-Mid DMRs, suggesting a potential overcompensation effect by DNMT3B in the epiblast on this specific DMR subtype. In comparison, *Dnmt3b-*depletion led to a substantial loss of average methylation levels in all DMR categories in the E6.5 epiblast and E×E, except for epiblast High-Mid DMRs (Figure 5A, Table S3) for which a compensatory mechanism provided high levels of methylation (wt: 76.9% vs *Dnmt3b*^−/−^: 71.6%). The compensatory mechanism on High-Mid DMRs in response to *Dnmt3b-*deficiency was ineffective in the E6.5 E×E. When we focused on promoter-TSS for each DMR categories, we observed again that loss of methylation for these regulatory regions was principally associated with *Dnmt3b*-deficiency (Figure 5B). The promoter-TSS regions associated with High-Mid DMRs retained relatively high methylation levels for either *Dnmt3a*^−/−^ or *Dnmt3b*^−/−^ epiblast samples, demonstrating a robust and compensatory *de novo* methylation mechanism. To further underline the impact of DNMT3A or DNMT3B on the *de novo* methylation of DMRs, we measured methylation levels for DMRs selected from our gene enrichment analyses (*Fgb*, *P2rx7*, *Pcyt2*, *Etnppl*, *Ralgds*, *Lrp5*) and other genomic segments covered by multiple tiles (*Cxxc1*, *Fcgrt*, *Lamp5*, *Mbd1*, *Lphn1*, *Pick1*, *Irf1*) (Figure S6 & S7) in *Dnmt3a-* or *Dnmt3b-*deficient epiblast and E×E. In line with our global observations, for most DMR-associated tiles, a *Dnmt3b-*deficiency in E6.5 epiblast or E×E caused a more severe loss of methylation compared to lack of *Dnmt3a.* However, for some regions in the E6.5 epiblast (e.g., *Pick1*, *Irf1*), *Dnmt3a*^−/−^ methylation levels were lower to those of *Dnmt3b*^−/−^. Overall, we show that of DNMT3A and DNMT3B participate in the establishment of the asymmetric methylation patterns associated with the various DMR categories in both embryonic and extraembryonic cells, with DNMT3B being the principal contributor in both cell lineages during the *de novo* reprogramming wave as it can compensate almost entirely for the absence of DNMT3A.

**Figure 5.**
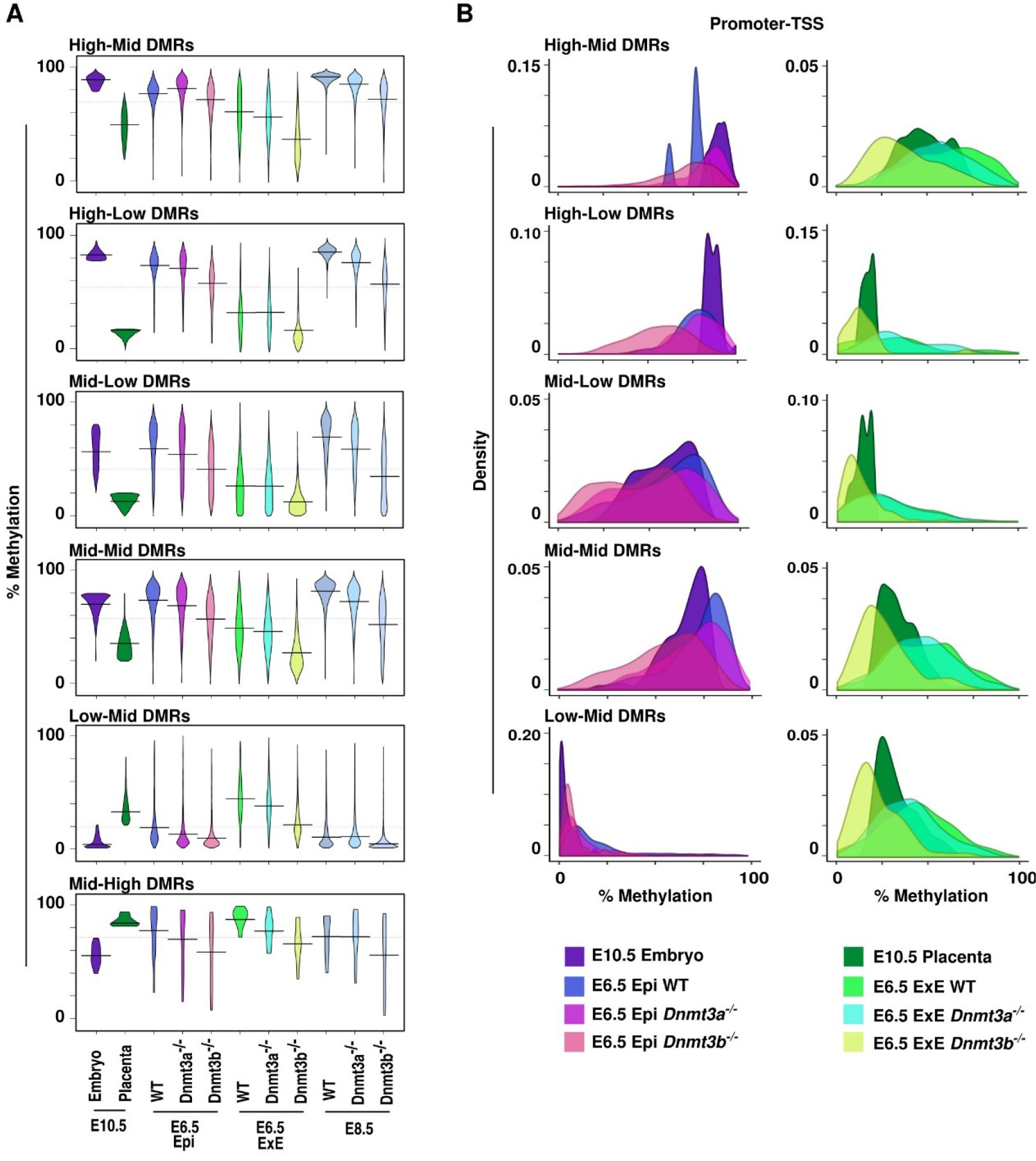
*Dnmt3a*- or *Dnmt3b*-deficiency alter proper establishment of DMR-associated patterns. **A)** Violin plot representing DNA methylation distribution and median values of tiles associated with the different DMR categories at E10.5, and their methylation levels in the different tissues and genotypes at E6.5 and E8.5. **B)** Promoter-TSS density graphs for E10.5 embryo and E6.5 epiblast (wt, *Dnmt3a*^−/−^and *Dnmt3b*^−/−^) (left panel) and for E10.5 placenta and E6.5 extraembryonic ectoderm (wt, *Dnmt3a*^−/−^and *Dnmt3b*^−/−^) (right panel). E: embryo, Epi: epiblast, E×E: extraembryonic ectoderm. See Table S3 for median and mean methylation values, and Table S4 for number of overlapping tiles analysed.

### Decline in Compensatory DNA Methylation Mechanisms in Response to Dnmt3a- or Dnmt3b- Deficiency

As embryonic development progresses from E6.5 to E8.5, our data suggest that lack of DNMT3A or DNMT3B further reduces global methylation levels of DMRs (High-Mid, High-Low, Mid-Low and Mid-Mid), evoking that absence of either enzymatic activity causes an additive and extended effect (Figure 5A-B). To further define the robustness in compensatory mechanism between DNMT3A or DNMT3B in the establishment and maintenance of DNA methylation on DMRs categories, we focused on High-Mid promoter-TSS- associated DMRs as they have the highest DNA methylation levels and require the most *de novo* methyltransferase activity.

Methylation differences of ≥20% between wild-type and *Dnmt3a*- or *Dnmt3b*-deficient samples were considered as regions showing substantial lack of compensation. In agreement with our results (Figure 5A-B), we observed that at E6.5, DNMT3B can compensate almost entirely for DNMT3A loss by maintaining methylation levels on most promoters-TSS associated tiles in the epiblast (410/437 = 93.8%) and E×E (558/602 = 92.7%), whereas DNMT3A can only partially alleviate the lack of DNMT3B in the epiblast (315/410 = 78.6%) and E×E (244/524 = 46.6%) (Figure 6A-B). As embryonic cell lineages development evolves between E6.5 and E8.5, global methylation levels increase on promoter-TSS of High-Mid DMRs. However, we detected a sharp decline in the compensatory mechanism to rescue the *de novo* and/or maintenance of methylation levels of promoter-TSS DMRs in the *Dnmt3a*- (524/627 = 80.9%) or *Dnmt3b-* (256/622 = 41.2%) deficient E8.5 embryos. This is further highlighted when we focus on a subgroup of 314 promoter-TSS associated tiles overlapping all of 9 data sets (Figure 6C), including gene promoters (e.g., *Asz1, Catsper1, Ccdc42, Dmrtb1, Piwil2, Rpl10l, Sox30, Sycp1, Tnp1, Ttll1, Zfn42*) related to our top enriched biological functions (i.e., piRNA, gamete generation and germ cells, meiotic nuclear division). Although E8.5 *Dnmt3a*- or *Dnmt3b*-deficient E×E data was not available, our data suggest that greater compensation failure would also be observed in this tissue. Thus, embryonic- placental DMRs need the combined action of DNMT3Aand DNMT3Bto both establish and maintain proper asymmetric levels during early development as compensatory DNA methylation mechanisms during the *de novo* wave of methylation decline overtime and fail to overcome methyltransferase shortage in later development stages.

**Figure 6.**
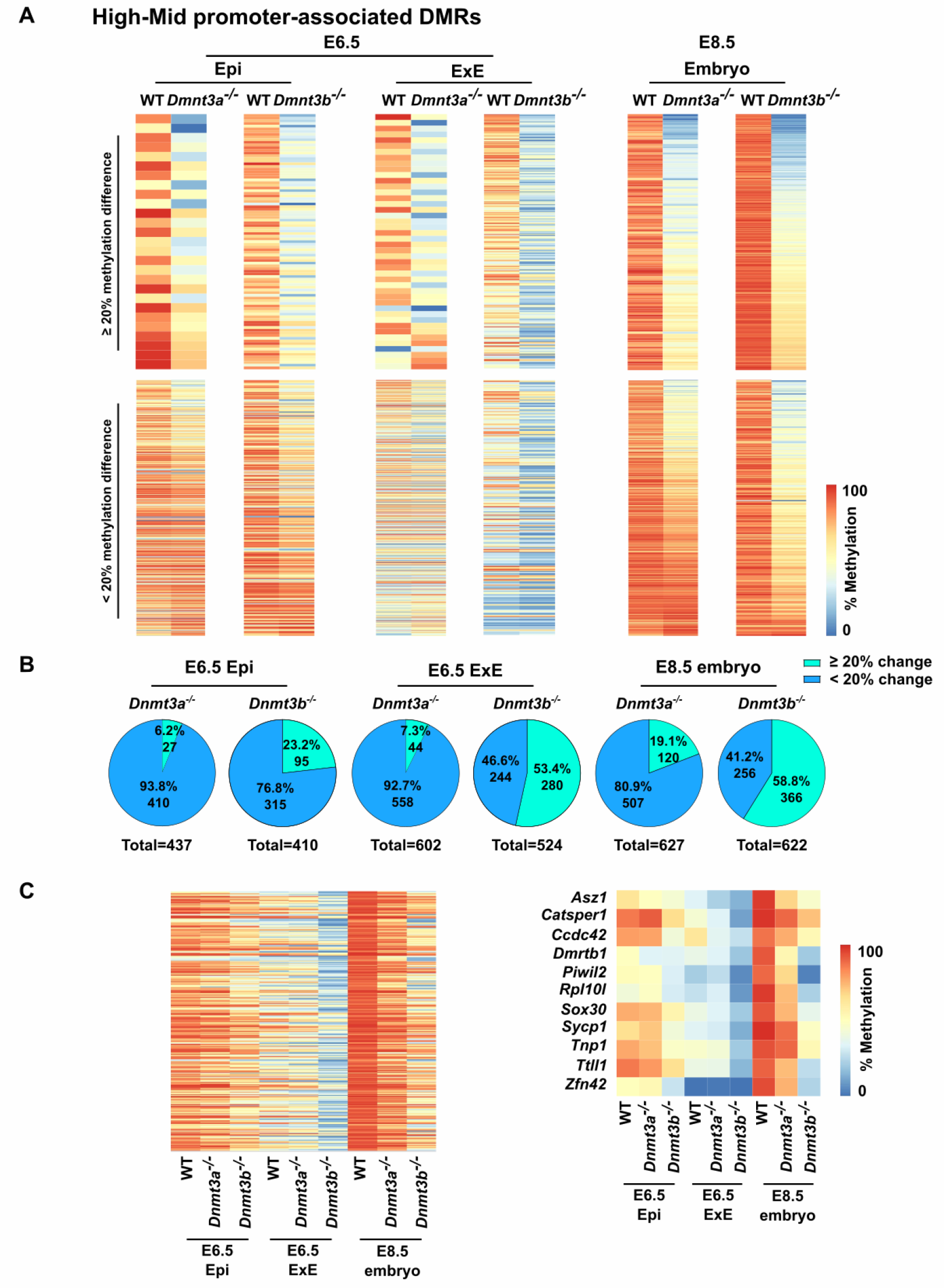
Developmental decline in compensatory DNA methylation mechanism in response to *Dnmt3a* - or *Dnmt3b*-deficiency at promoter-TSS associated with High-Mid DMRs. **A)** Heatmaps comparing the DNA methylation profiles of wt vs *Dnmt3a*^−/−^ or wt vs *Dnmt3b*^−/−^ for E6.5 epiblast, E6.5 extraembryonic ectoderm or E8.5 embryo in 100bp tiles overlapping promoter-associated DMRs that are highly methylated in the embryo (≥80%) and mildly methylated in the placenta (≥20 and <80%). Upper panel represents regions with a difference of methylation of at least 20% between wt and *Dnmt3a*^−/−^ or wt and *Dnmt3b*^−/−^. Lower panel represents stable regions (lower than 20% difference in methylation). **B)** Pie charts representing DMR numbers for each comparison. **C)** Heatmap of the methylation levels of tiles commonly represented in all 9 public data samples that overlap with promoters-associated High-Mid DMRs (n = 314 tiles) (left panel). Heatmap of the methylation level of gene associated with piRNA, gamete generation and germ cells or meiotic nuclear division that are covered in all 9 public datasets (right panel).

## DISCUSSION

With recent breakthroughs in high-throughput sequencing, we now have a better understanding of the dynamic of DNA methylation erasure occurring in early cleavage stage embryos following fertilization. However, the discrepancies in acquisition of genome-wide DNA methylation patterns in early post-implantation embryonic and extraembryonic cell lineages remain to be methodically delineated. To address this issue and further our understanding of the DNA methylation asymmetry that guides the developmental trajectory of embryonic and extraembryonic cell lineages, we established genome-wide DNA methylation profiles of mouse embryo and placenta at mid-gestation, and analyzed various publicly available developmental stage specific embryo and placenta DNA methylation datasets. Using this strategy, we uncovered that 45% of the genomic regions analyzed differ in DNA methylation status (≥20%) between mid-gestation embryo and placenta, and that these DMRs can be further divided into categories based on their levels of DNA methylation (Low; <20%, Mid; ≥20 to <80%, High; ≥80%) in the embryo and placenta. We show that the embryo and placenta acquire specific DMR categories during the early stage of the *de novo* DNA methylation wave, and that these DMRs persist throughout prenatal development, as well as into somatic adult tissues. Furthermore, we show that *Dnmt3b* primarily drives the divergence in DNA methylation levels associated with these specific DMRs and that *de novo* methyltransferase activity of *Dnmt3b* can almost entirely compensate for lack of *Dnmt3a* in the asymmetric establishment of embryonic and placental DMRs, but that *Dnmt3a* can only partially compensate the absence of *Dnmt3b*. However, with developmental progression, this compensatory DNA methylation mechanism becomes less effective.

Our results indicate that the kinetics of DNA methylation acquisition leading to specific embryo-placenta DMR categories is not a stepwise process occurring throughout cell fate decisions and patterning of embryonic and extraembryonic lineages, but a prompt progression in the early post-implanted conceptus. These results are in line with studies reflecting that the initiation of asymmetric DNA methylation levels begins within the trophoblast and the inner cell mass of the blastocyst (Santos et al., 2002), and becomes highly evident by the time the epiblast acquires its initial global DNA methylation patterns at E6.5 during the *de novo* reprogramming wave (Auclair et al. 2014, Smith et al. 2017). Our results show that the acquisition period between E4.5 and E6.5 is particularly key to establishing asymmetric DNA methylation patterns associated with mid-gestation embryo-placenta DMR classes, and that these DMR associated-patterns are long-lasting across stages of prenatal development. Since we studied cell populations derived from embryonic and placental tissues, we cannot dismiss the prevalence of cell-to-cell heterogeneity in the acquisition kinetics of specific DMR patterns. Despite this concern, our results indicate that embryonic cells have a large body of DMRs with High-levels (≥80%) and Low-levels (<20%) of methylation that remain static across development, as well as in somatic cell types (High-Mid & High-Low DMRs), whereas compared to the embryo, the placental DMRs have overall lower methylation levels and are more broadly distributed amongst methylation levels, which has also been shown in other studies where genome-wide DNA methylation profiles of placental cells were compared to other tissues and specific cell types (Schroeder et al., 2013; Chatterjee et al., 2016; Smith et al., 2017). Lower methylation levels in the placenta have been associated with reduced *de novo* methyltransferase activity in the E×E during the *de novo* reprogramming wave (Fulka et al., 2004; Santos et al., 2002). Nevertheless, we observe methylation level peaks for all DMR categories within the E×E at E6.5 before levels stabilize at E10.5. As of now, the implication of these methylation level peaks on future regulation mechanisms and methylation profiles is unknown. Moreover, it remains to be determined whether global reduction in DNA methylation marks on DMRs between the E6.5 and E10.5 extraembryonic cell lineages is stochastic or targeted, and whether it occurs through passive or active mechanisms.

We also observed an enrichment of retrotransposons (i.e., LINE, SINE, LTR) in DMRs especially in those with High-levels of methylation in the embryo and lower-levels in the placenta, whereas DMRs with Low-levels of methylation in the embryo and higher levels in the placenta are preferentially outside retrotransposons-associated sequences. Earlier findings of Chapman et *al.* (Chapman, Forrester, Sanford, Hastie, & Rossant, 1984) revealed that repeat regions in the placenta appear to lack tight control of their methylation patterns, perhaps indicating that maintaining methylation, and therefore repression of these elements for genome stability and integrity, is not critical given the relatively short lifespan of this organ. However, we do observe Mid-range methylation levels (20-80%) in the placenta for a substantial portion of DMRs that are associated with retrotransposons, revealing that specific genomic regions associated with repetitive elements do need tight regulation in extraembryonic cell lineages for proper development. These results are in line with findings exposing that the deletion of genome-defense gene *Tex19.1* leads to the de-repression of LINE1 and compromises placental development, suggesting that disparities between retrotransposon suppression and genome-defense mechanisms might contribute to placenta dysfunction and disease (Reichmann et al., 2013).

DNMT3A and DNMT3B are required to establish proper methylation profiles on the embryonic genome during the *de novo* reprogramming wave, as both methyltransferase enzymes have redundant, but also specific functions. However, the activity of DNMT3B is the main contributor in the acquisition of profiles in epiblast cells, and especially commands methylation on CGIs associated with developmental genes (Auclair et al., 2014). Auclair et *al.* highlighted that in the absence of DNMT3B, DNMT3A was not able to counterbalance, leading to the loss of promoter-CGI methylation and gain of expression of germline genes (e.g., *Sycp1*, *Sycp2*, *Mael*, *Rpl10l*, *Dmrtb1*) in somatic cells of the embryo. Here, we show that the promoter of these germline genes, associated with meiotic and piRNA processes as well as other genes with similar biological functions, are highly enriched in High-Mid DMRs. We also reveal that for this particular set of High-Mid DMRs, mainly deprived of CGIs, DNMT3A is able to substantially counterbalance the lack of DNMT3B in the E6.5 epiblast and establish global DNA methylation levels, whereas in the E×E, this compensatory mechanism is rather inefficient. Globally, we show that DNMT3B is much more potent at compensating than DNMT3A in both the epiblast and E×E for all DMR categories, indicating that DNMT3B is the main *de novo* enzyme driving asymmetric DNA methylation patterns between the embryo and placenta. Although compensatory mechanisms have been observed in *Dnmt3a* or *Dnmt3b*-depleted conceptuses, we still do not fully understand the process as *Dnmt3a* and *Dnmt3b* have cell lineage specific expression during the peri-implantation developmental period. As development progresses, we observed that the compensation mechanism in *Dnmt3a* or *Dnmt3b*-deficient embryos remains apparent at E8.5 on most DMR categories, but is less effective as the methylation gaps increase compared to wild-type. This correlates with a developmental period where DNMT3A is now the main *de novo* methyltransferase enzyme in both the embryo and placenta (Okano et al., 1999; Watanabe et al., 2002), and with a compensatory activity of DNMT3A being less effective. This suggests that DNMT3B activity is critical to ensure proper establishment of DNA methylation asymmetry between the embryonic and extraembryonic cell lineages during the *de novo* reprogramming wave, but that DNMT3A is required during the developmental progression to safeguard methylation levels on DMR categories.

## CONCLUSION

We demonstrate that asymmetry between embryo and placenta DNA methylation patterns occurs rapidly during *de novo* acquisition of methylation marks in the early post-implanted conceptus, and that these patterns are long-lasting across subtypes of DMRs. We also reveal that at the peri-implantation stages, *de novo* methyltransferase activity of DNMT3B is the main provider of asymmetric methylation marks on DMRs, and that it largely compensates the lack of DNMT3A in the epiblast and E×E. However, as development progresses, DNMT3A becomes the principal *de novo* methyltransferase by mid-gestation, and DNMT3B methyltransferase activity is less effective at promoting compensation. These results further underline why embryos developing without DNMT3B have severe DNA methylation defects and die at mid-gestation, whereas those without DNMT3A only die postnatally. Further investigation is required to determine the molecular mechanisms controlling the precise *de novo* acquisition of long-lasting methylation marks on specific DMR subtypes in the embryonic and extraembryonic cell lineages, and how errors in this process could lead to abnormal development and diseases.

## ACKNOWLEDGMENTS

We thank the McGraw lab for critical comments and suggestions, as well as Elizabeth Maurice- Elder for editing.

## DATA AVAILABILITY

The data from this study have been submitted to the Gene Expression Omnibus (#GSE95610). Reviewer link: https://www.ncbi.nlm.nih.gov/geo/query/acc.cgi?token=ghsnmqoctlgjlcj&acc=GSE95610

---

**Table S1. Annotation and DNA methylation values associated with DMRs.**

**Table S2. DNA methylation levels for DMR categorises across development.**

**Table S3. DNA methylation levels for DMR categories in various *Dnmt* knockout mouse models.**

**Table S4. Number of 100bp tiles in each category of public data that overlap with Embryo-Placenta DMRs (related to Figure 4; 5A; S4; S5)**

