## Supplemental Figures for "Developmental Genome-Wide DNA Methylation Asymmetry Between Mouse Placenta and Embryo"

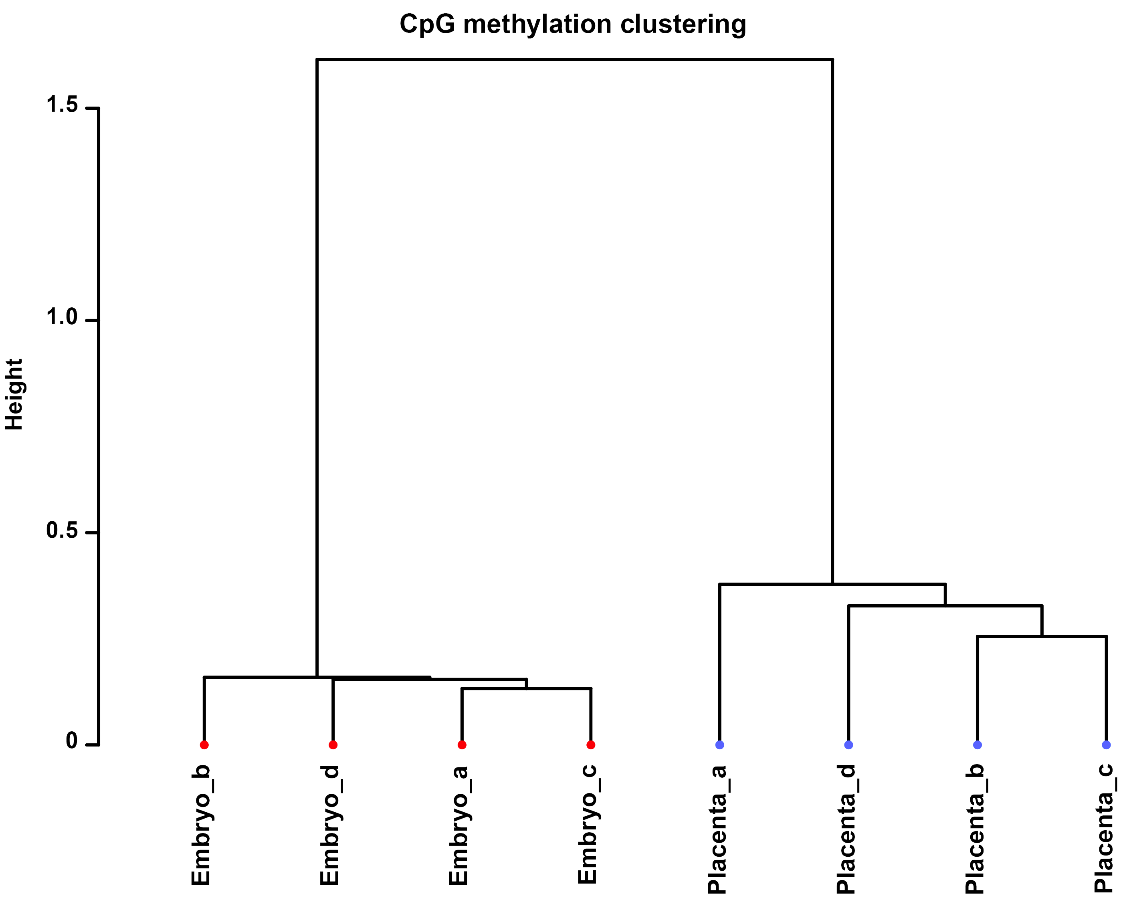

**Figure S1. Hierarchical dendrogram showing clustering of E10.5 embryo and placenta samples according to genome-wide DNA methylation profiles. Samples *a, c*: females; Samples *b*, *d*: males.**

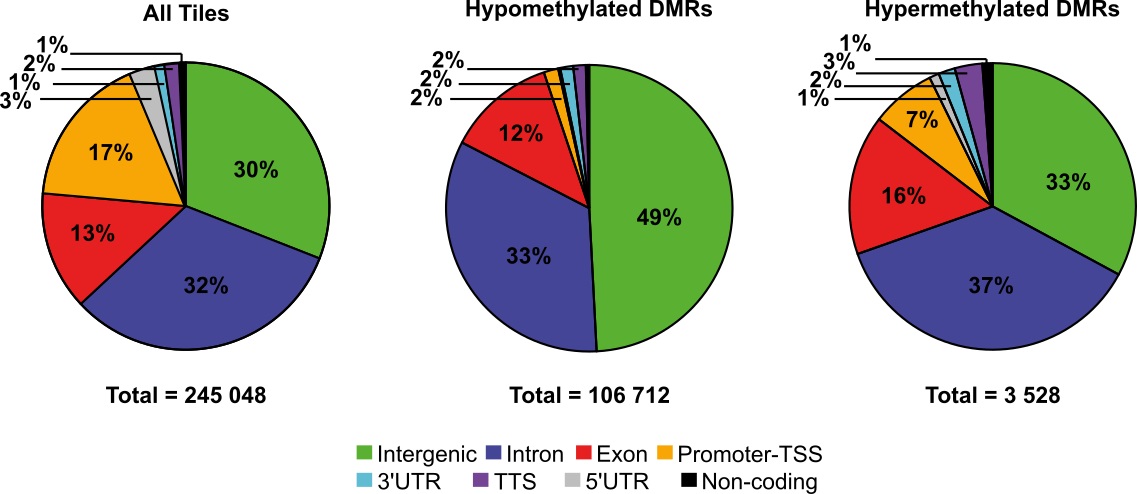

**Figure S2. Pie charts representing genomic annotation of Hypo- and Hyper-DMRs found between E10.5 embryo and placenta.** Proportion of all sequenced tiles (All-tiles; n=245 048) common to embryo and placenta, Hypo-DMRs (n=106 712) and Hyper-DMRs (n=3 528), found in intergenic and genic regions (exons, introns, promoters-TSS, 3′ and 5′ untranslated regions, and transcription termination sites, non-coding).

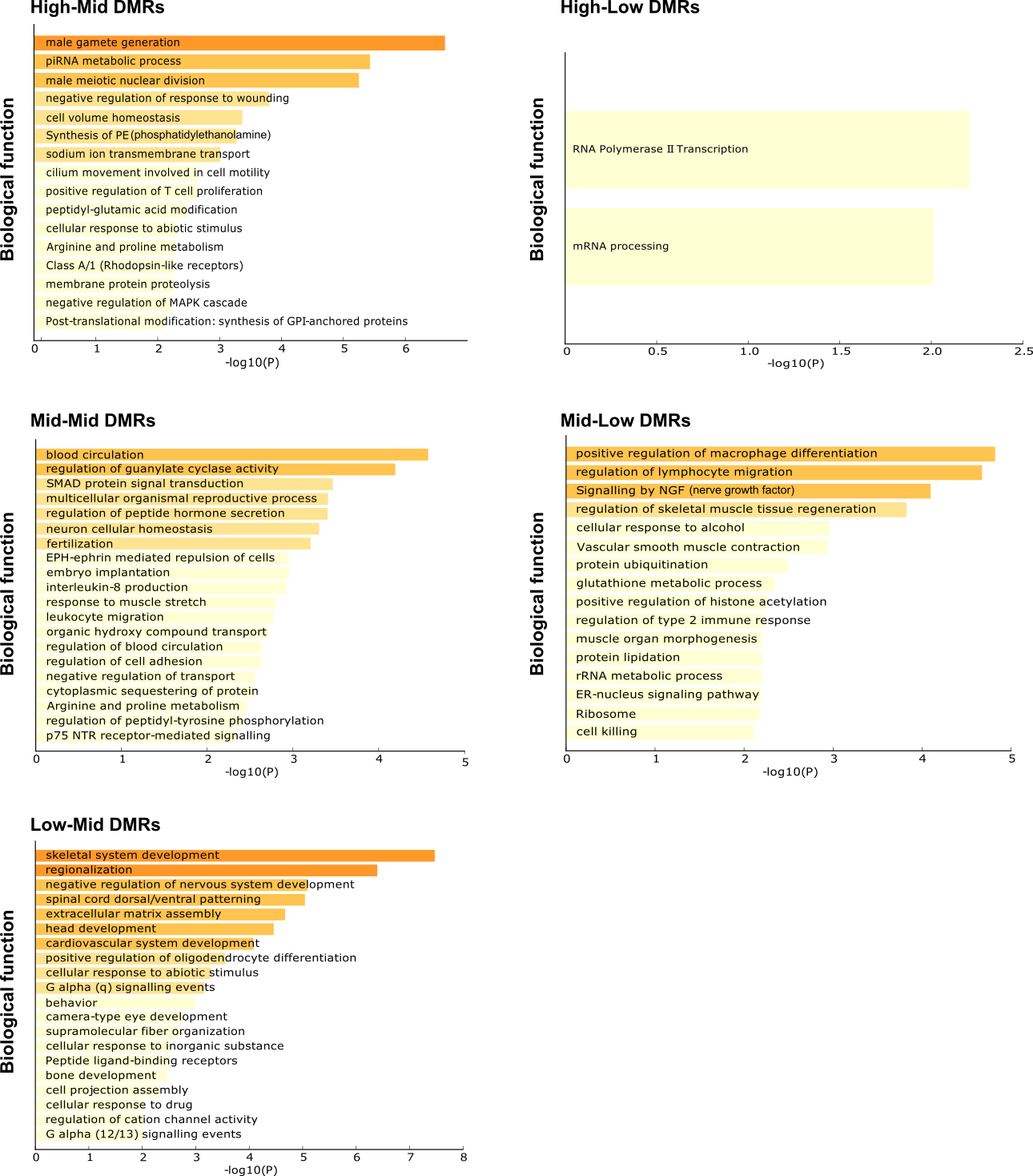

**Figure S3.** **Summary of biological functions associated with E10.5 embryo-placenta DMRs subtypes.** Number of tiles in promoter-TSS regions used as input for each methylation category: High-Low: 38; High-Mid: 749; Mid-Lo: 477; Mid-Mid: 509 and Low-Mid: 235. No enrichment in biological functions found for Mid-High: 1.

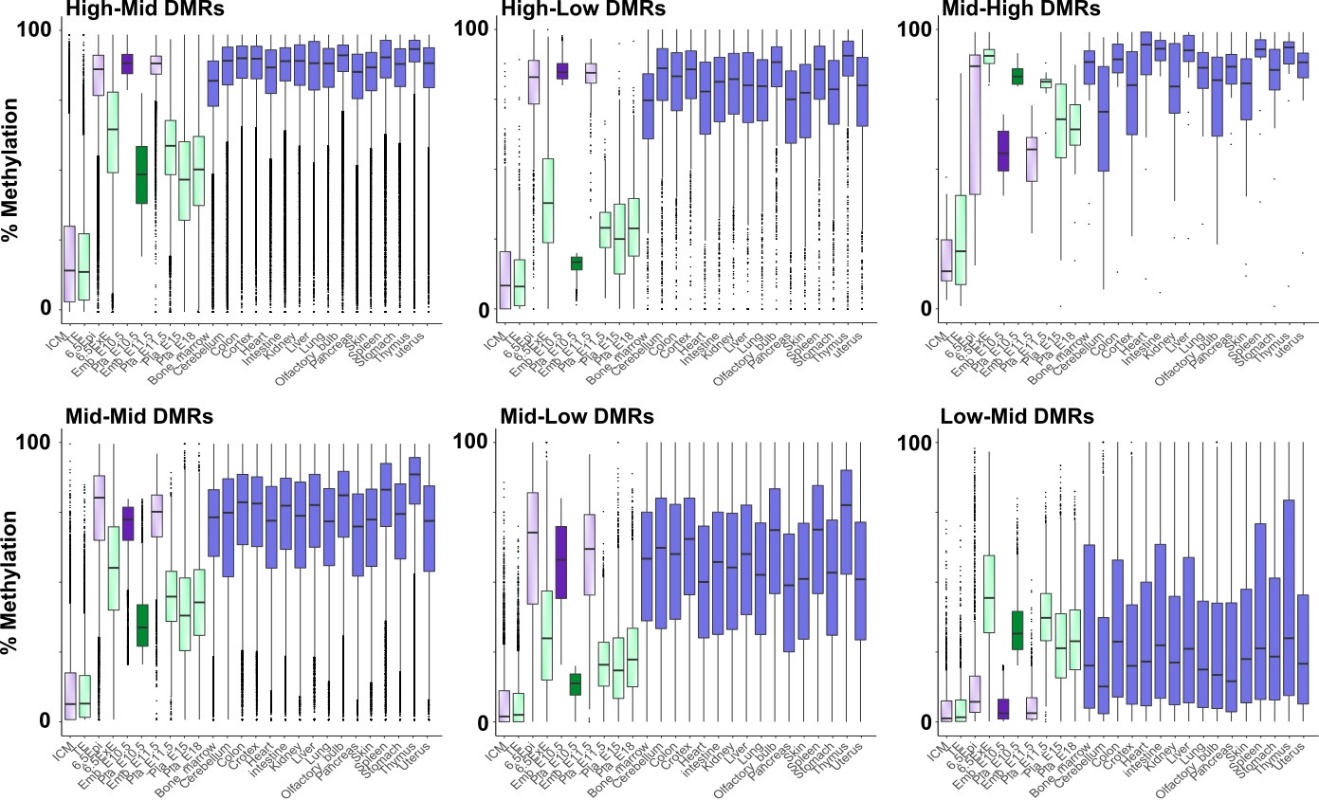

**Figure S4. Dynamics of DMR regions during embryonic development and in adult tissues.** Box-plots showing DNA methylation distribution and median values of overlapping DMRs to publicly available methylation data at different developmental stages and in different adult tissues (Smith et al., 2014, Whidden et al., 2016, Hon et al., 2013, Decato et al., 2017). ICM: inner cell mass, TE: trophectoderm, Epi: epiblast, ExE: extraembryonic ectoderm, Emb: embryo, Pla: placenta. See Table S2 for median and mean methylation values.

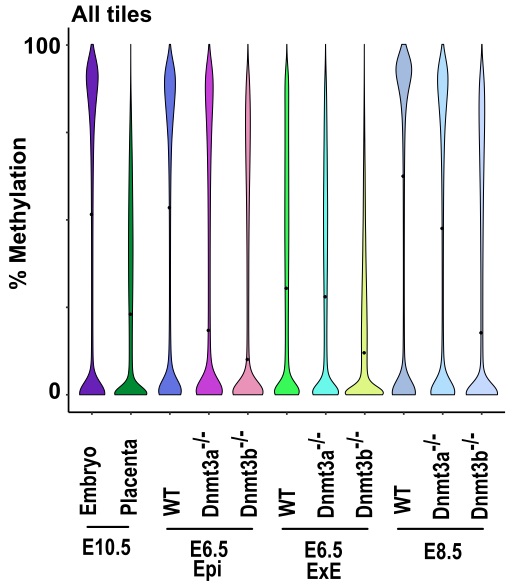

**Figure S5. Global DNA methylation profiles in *Dnmt3a^-/-^* and *Dnmt3b^-/-^* samples.** Violin plot showing DNA methylation distribution and median values of *Dnmt3a* or *Dnmt3b* knockout mice in E6.5 epiblast (Epi), E6.5 extraembryonic ectoderm (ExE) and E8.5 embryos in 100bp tiles corresponding to the 245 048 analyzed tiles. See Table S3 for median and mean methylation values.

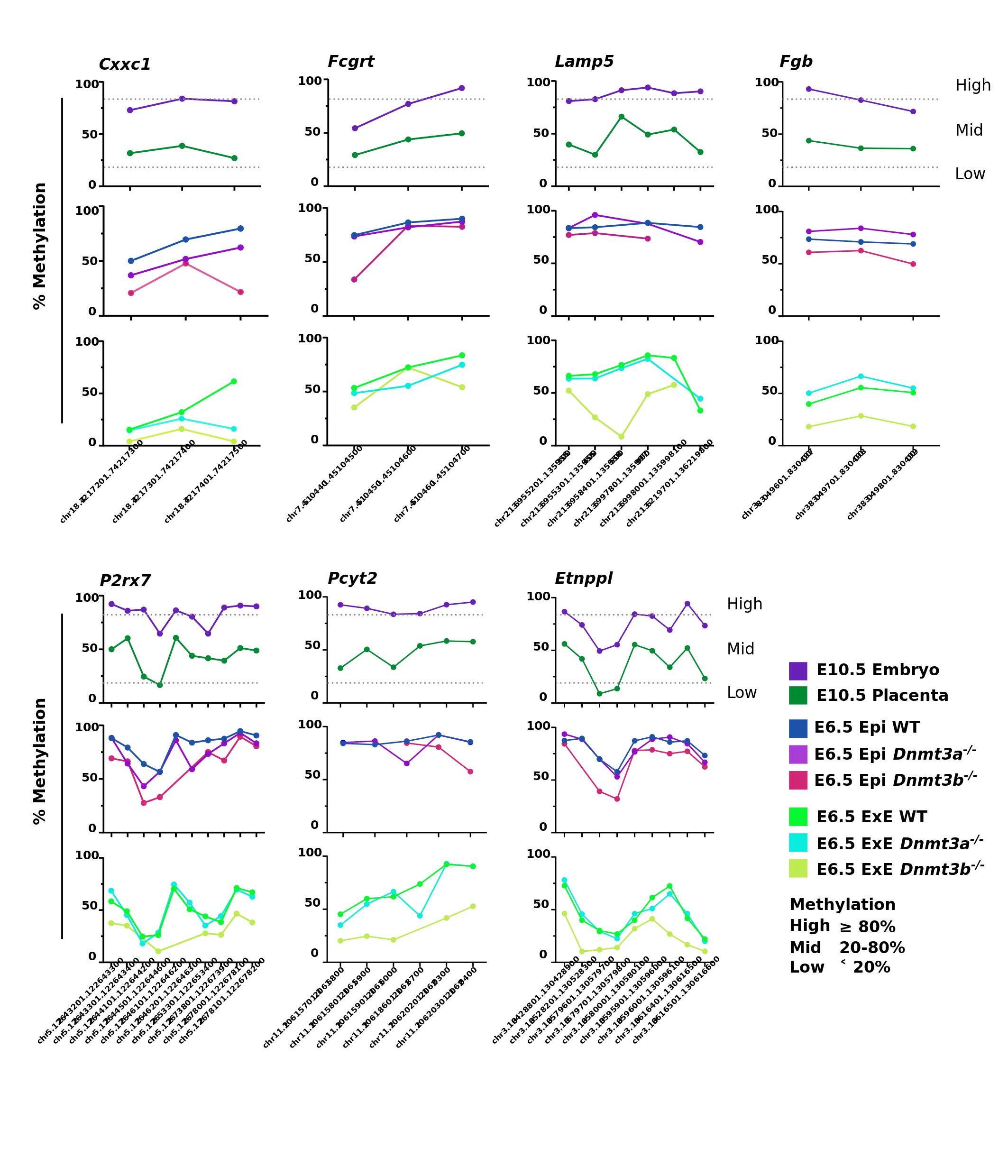

**Figure S6.** ***Dnmt3a*- or *Dnmt3b*-deficiency alter proper establishment of DNA methylation associated with various DMR categories within gene promoter-TSS.** DNA methylation average (%) per tile in *Dnmt3a^-/-^* and *Dnmt3b^-/-^* E6.5 epiblast and extraembryonic tissues for E10.5 embryo-placenta associated DMRs. *Lrp5* (activation of *Wnt* signaling, important role in development processes); *Fgb* (encode for beta component of fibrinogen); *P2rx7* (ATP receptor); *Pcyt2* (role in the biosynthesis of phospholipid phosphatidylethanolamine); *Etnppl* (role in the biosynthesis of glycerophospholipid and metabolism); *Ralgds* (role in signaling processes).

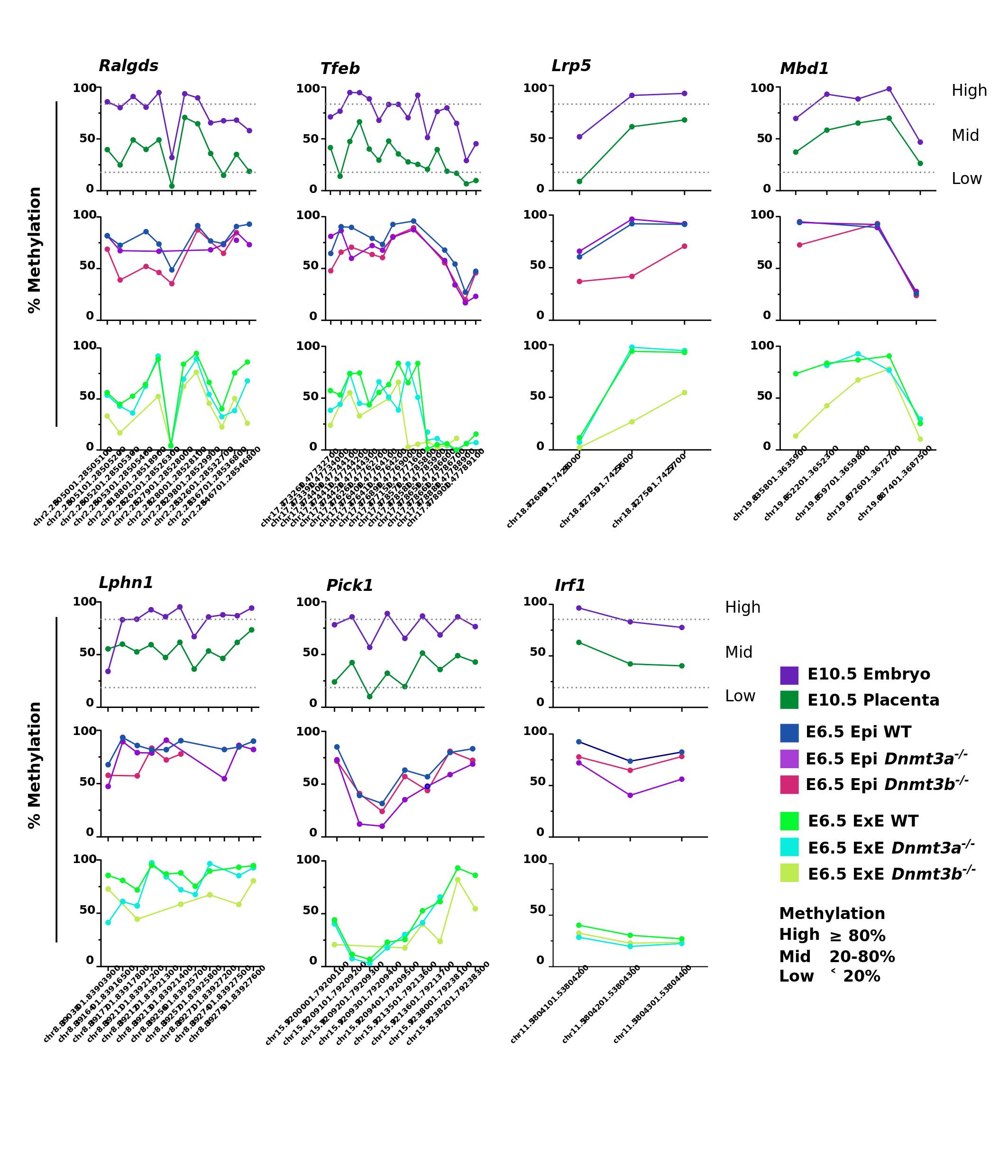

**Figure S7.** ***Dnmt3a*- or *Dnmt3b*-deficiency alter proper establishment of DNA methylation associated with various DMR categories within genic regions.** DNA methylation average (%) per tile in *Dnmt3a^-/-^* and *Dnmt3b^-/-^* E6.5 epiblast and extraembryonic tissues for E10.5 embryo-placenta associated DMRs. *Cxxc1* (role in regulation of gene expression and development); *Fcgrt1* (transfers IgG from mother to fetus across the placenta); *Lamp5* (role in synaptic plasticity in some GABAergic neurons); *Mbd1* (transcriptional repressor); *Lphn1* (role in cell adhesion and signal transduction); *Pick1* (role in synaptic plasticity and regulation of astrocyte morphology); *Tfeb* (role in the regulation of lysosomal genes and autophagy); *Irf1* (regulation of cellular response, including *IFN* and *IFN*-inducible-genes).
